# When monkeys meet an ANYmal robot in the wild

**DOI:** 10.1101/2024.08.13.607714

**Authors:** Charlotte Canteloup, Joonho Lee, Samuel Zimmermann, Morgane Alvino, Markus Montenegro, Marco Hutter, Erica van de Waal

**Affiliations:** Department of Ecology and Evolution, University of Lausanne, Switzerland; INKAWU Vervet Project, KwaZulu-Natal, South Africa; The Sense Innovation and Research Center, Lausanne and Sion, Switzerland; Laboratory of Cognitive and Adaptive Neurosciences, CNRS - UMR 7364, University of Strasbourg, Strasbourg, France; Robotic Systems Lab, ETH Zürich, Switzerland; Neuromeka Co., Ltd. Korea

**Keywords:** animal-robot interaction, primate, ethorobotics, behavior, field experiment

## Abstract

Animal-robot interaction studies have been of increasing interest in research, but most of these studies have involved robots interacting with insects, birds, and frogs in laboratory settings. To date, only two studies used non-human primates and no behavioral study has tested the social integration of a robot in a group of wild primates. To fill this gap, we studied the interactions between the quadruped ANYmal robot and a group of 37 wild vervet monkeys in South Africa. The ANYmal robot is a remote-controlled sheep-sized robot with an open box of food on its back. We gradually introduced the robot to the monkeys following five different steps over 6 days for a total exposition time of about 10h. The monkeys habituated to the robot very quickly with six individuals eating the food in the robot’s box from the second day. A few individuals, mostly juveniles, emitted alarm calls towards the robot. In total, seven individuals, high rankers, spent time near the robot and 21 monkeys approached the robot from a greater distance. High rankers displayed significantly more vigilant and self-centered behaviors and they, with females and juveniles, ate more food in the robot’s box compared to low rankers, males, and adults conversely. This study offers exciting perspectives on the phenomena of social acceptance of machines in mammalian societies and the automation of field data collection.

## Introduction

From food vending machines to domestic, military, and industrial robots, including artificial intelligence, robots have an exponentially growing important place in our society. In recent decades, robots have also been increasingly used in animal studies^1,2^. We define here a robot as “a machine that is able to interact physically with its environment and perform some sequence of behaviors, either autonomously or by remote control”^1^. The first studies date back to the fifties when some researchers such as Tinbergen^3^ used manually manipulated dummies and decoys, to study the social behaviors of three-spined sticklebacks and gulls. Remote-controlled robots that perform a pre-programmed sequence of behaviors have subsequently been used in various animal-robot interaction studies. On the one hand, biomimetic robots mimicking the studied species have been designed to test whether communicative signals displayed by the robotic model could elicit a behavioral response in real animals^4,5^. For example, some researchers have investigated what kind of communicative signals are considered by squirrels to communicate an alarm^6^, by frogs to defend their territory^7^ and during courtship^8^ , or whether starlings responded to the orienting behavior of a robotic conspecific^9^. Biomimetic robots have also been used to test whether locusts can use social information provided by a robotic demonstrator to avoid predators^10^ or whether a robotic fish can recruit real fishes from a refuge and initiate new swimming direction^11^. These studies suggest that biomimetic robots can be perceived by animals as conspecifics. On the other hand, other types of non-biomimetic robots that do not perfectly match the species under study, have been used to test whether animals can recognize them as social partners. Authors found that dogs responded more to a furry dog-like AIBO robot compared to a remote-controlled car^12^, but less to a pointing humanoid robot than to a human^13^; both dogs and cats discriminated between animate and inanimate unidentified moving objects^14^, but without any evidence that they developed individual recognition of them^15^. One study reported that chicks raised with a robot improved their spatial abilities^16^. Interestingly, rats learnt from a rat-sized robot a lever-pushing task to obtain food^17^ and they even behaved pro-socially towards a robotic rat, by releasing it from restraints, especially if the robot had previously been helpful to them compared to an unhelpful robot^18^. In primates, marmosets have been reported to attribute goals to a small four-legged robot but not to a moving box^19^ and chimpanzees were particularly sensitive to a doll-like robot that reproduced their behavior, from which they even requested social responses^20^. These studies suggest that animals can perceive unfamiliar robots as animate to some extent based on some lifelike cues, such as a body with a head and legs, biological motion, and self-propulsion. With the development of technology, some robots have been designed with an autonomous mode, so that they can interact with their environment, learn, and even adapt, leading some scientists to claim for ‘mixed societies of animals and robots’^21,2^. Mobile robots capable of detecting obstacles, adjusting their trajectories, and controlling certain group behaviors, have been introduced in the lab to precocial chicks. The chicks followed the robots, aggregated with them, and developed an attachment to them through the learning process of filial imprinting, showing distress after separation^21,22^. Autonomous robots have been notably used to investigate animal collective behaviors. Such robots have been socially integrated into cockroaches’ groups, leading groups’ decisions to move to shelters^23^. In the same vein, autonomous robots have been successfully developed to manoeuvre flock of ducks^24^, to investigate self-organisation in ant colonies^25^ and to create biohybrid systems by coordinating the collective behaviors of honeybees and zebrafishes using socially integrated robots^26^.

Most of the animal-robot interaction studies took place in laboratories with insects^10,23,25,26^, fishes^2,11^, frogs^8^, birds^5,9,16,21,22,27^ and, within mammals, with rats^17,18^ and dogs^13–15,28^. Only two studies have been conducted so far with primates, both in captivity, one with marmosets^19^ and one with chimpanzees^20^. Comparatively, far fewer studies have been conducted in the wild, with some species of insects^29^, frogs^7^, crocodiles and lizards^30^, birds^4,31^ and squirrels^6^. To our knowledge, no animal-robot interaction studies have been done with wild primates. To fill this gap, we introduced an ANYmal robot^32–34^ (Fig. 1) to a group of 37 vervet monkeys living in their natural habitat in South Africa. The ANYmal robot is a highly mobile and sophisticated four-legged robot designed for autonomous operation in harsh environment. The use of artificial agents with unfamiliar embodiment allows for high flexibility of motion without the influence of the familiar physical appearance, thus providing high control and repeatability. We report here a first study that aimed at investigating the reaction of the monkeys to the robot and their interaction in the field.

**Figure 1.**
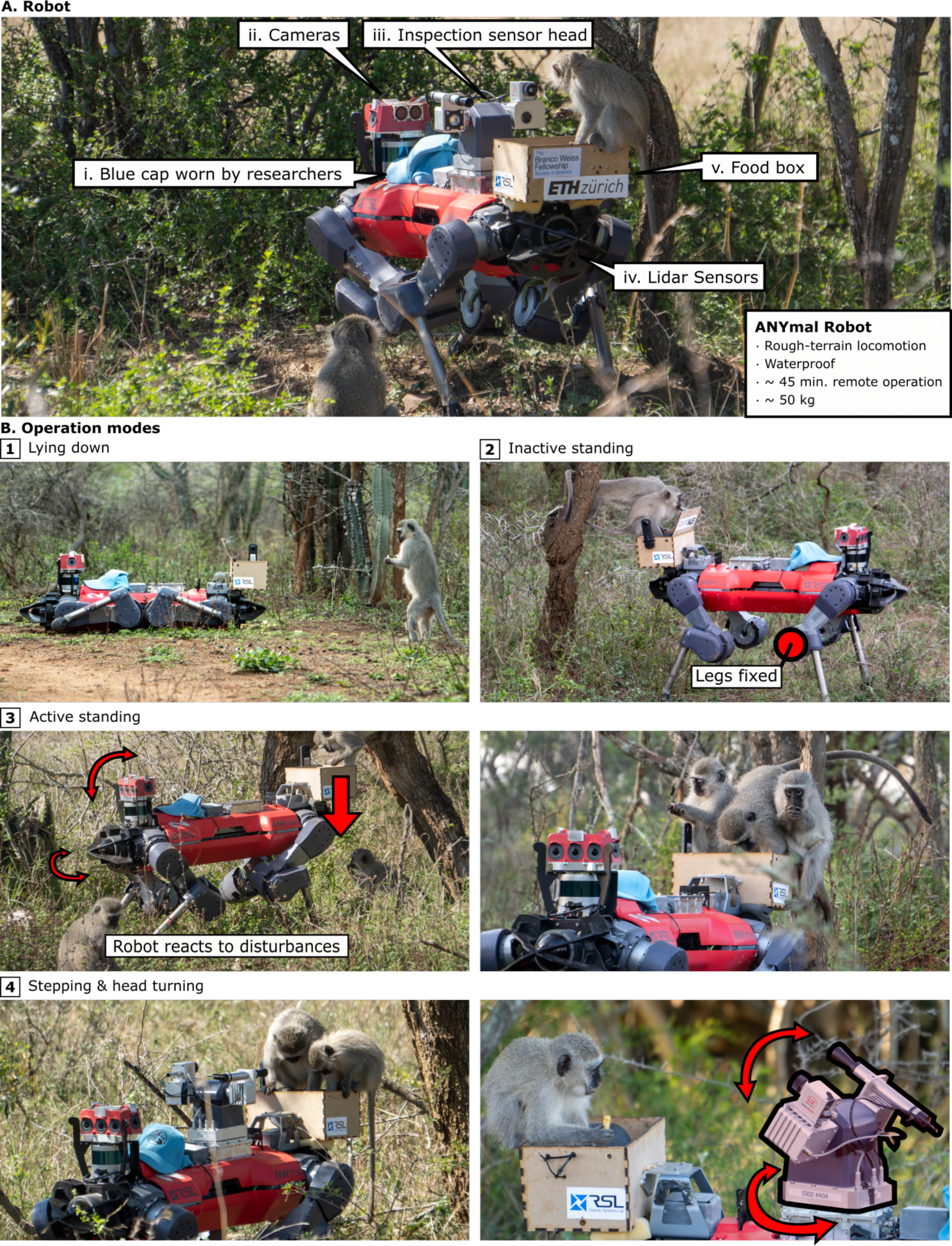
A) The ANYmal robot used in the experiment. B) Operation modes of the robot. We started from a static (1) lying down mode and gradually added more motions (3, 4). Picture credit: Joonho Lee

### Presentation steps of the ANYmal robot to the monkeys

The field experiment consisted of introducing an ANYmal robot (Fig. 1) to the group of wild vervet monkeys for one week. Because the robot was big, compared to the monkeys, we gradually introduced it to the group of monkeys following four different steps to avoid frightening them (Table 1; Table SI_1; Fig. 1B-4).

**Table 1.** Summary table of the experiment by step; robot position; food position; date; time (start time of the experiment); session number; experiment duration (in minutes: seconds); number of agonistic behaviors displayed by monkeys towards the robot; number of alarm calls emitted by monkeys in response to the robot; number of individuals eating food on the ground; [Numbers in brackets represent the individuals that ate food leftovers on the ground]; number of individuals eating food inside the box on the robot’s back; number of individuals present in contact and within arm length distance to the robot during scans; number of individuals present within more than arm length distance and up to 10m to the robot during scans.

| Step | Robot position | Food location | Date | Time | Session number | Experiment duration | Agonistic behaviours | Alarm calls | Eat food ground | Eat food box | contact-arm length distance | > arm-length-10m distance |
| --- | --- | --- | --- | --- | --- | --- | --- | --- | --- | --- | --- | --- |
| 1 | lying down | ground + box | 07/10/2021 | 11:40 | 1 | 24 | 0 | 0 | 0 | 0 | 0 | 0 |
| 1 | lying down | ground + box | 07/10/2021 | 16:31 | 2 | 19 | 0 | 3 | 6 | 0 | 3 | 10 |
| 1 | lying down | ground + box | 07/10/2021 | 17:08 | 3 | 12 | 1 | 0 | 3 | 0 | 3 | 5 |
| 1 | lying down | ground + box | 07/10/2021 | 17:27 | 4 | 37 | 0 | 2 | 3 | 0 | 0 | 4 |
| 1 | lying down | ground + box | 08/10/2021 | 05:32 | 5 | 53 | 6 | 2 | 11 | 6 | 6 | 12 |
| 1 | lying down | ground + box | 08/10/2021 | 06:30 | 6 | 26 | 0 | 0 | 6 | 0 | 0 | 7 |
| 2 | inactive standing | ground + box | 08/10/2021 | 07:17 | 7 | 40 | 4 | 0 | 3 | 0 | 0 | 5 |
| 2 | inactive standing | ground + box | 09/10/2021 | 05:39 | 8 | 57 | 9 | 0 | 2 | 6 | 6 | 11 |
| 2 | inactive standing | box | 09/10/2021 | 06:55 | 9 | 19 | 0 | 0 | 0 | 0 | 0 | 0 |
| 2 | inactive standing | box | 09/10/2021 | 07:45 | 10 | 38 | 0 | 0 | 0 | 6 | 6 | 8 |
| 3 | active standing | box | 09/10/2021 | 16:51 | 11 | 26 | 3 | 0 | 1 | 5 | 4 | 6 |
| 3 | active standing | box | 10/10/2021 | 16:59 | 12 | 47 | 37 | 1 | - | 5 | 4 | 9 |
| 4 | moving & head turning | box | 11/10/2021 | 08:41 | 13 | 66 | 0 | 0 | [3] | 6 | 5 | 14 |
| 4 | moving & head turning | box | 11/10/2021 | 16:25 | 14 | 60 | 6 | 0 | [4] | 5 | 5 | 8 |
| 4 | moving & head turning | box | 12/10/2021 | 05:40 | 15 | 81 | 14 | 4 | [3] | 6 | 7 | 12 |

We experimented with four different operation modes of the robot: lying down mode, inactive standing mode, active standing mode, and stepping/head-turning mode. In the lying down mode, the robot was inactive. In the inactive standing mode, the robot stood on its four legs with the leg joints fixed. In the active standing mode, the robot stood still but reacted to external disturbances and moved its body depending on the surroundings. In the stepping/head-turning mode, the robot stepped or turned its inspection head (Fig.1B-4) following the operator’s commands.

During the first step, the robot was lying down, with food inside the box and spread on the ground around the robot (Fig.1B-1). In the second step, the robot was in inactive standing mode. In this mode, the robot did not react to the monkeys. Food was placed inside the box and spread on the ground (Fig.1B-2). Although still cautious, the monkeys began to approach and touch the robot.

In the third step, the robot was in active standing mode (Fig.1B-3), moving its body slightly in response to the approaching monkeys and the additional weight. This movement made the monkeys more vigilant than in the previous step. There was food only inside the box. In the final step, the robot was in stepping/turning mode, moving its body slightly and occasionally turning its inspection head 360°. There was only food inside the box (Fig. 1B-4).

On the first day, we brought the robot to the field in the late morning when the group of monkeys was more than 100m away across the river, but no individual got closer to the robot. The second session consisted of the first encounter between the robot and the monkeys that occurred later in the same day in another part of the monkeys’ territory. Two minutes into the session, one adult female (Guat) approached the robot that was lying down on the ground within one meter, stood-up bipedally for few seconds while looking inside the box on the robot’s back, and immediately left. In total, six individuals from the dominant ‘G’ matriline (Table_SI_1) approached and started to eat the food on the ground in front of the robot during this session. On the second day, six individuals started to eat food inside the robot’s box (Table 1; Table SI_1). Overall and across the experiments, eight individuals emitted 12 alarm calls towards the robot (Table 1; Table SI_1). A total of four individuals emitted 80 agonistic behaviors, mostly head bobbing towards the robot. An individual (Gri) jumped on the robot for the first time on the third day (session 10; step 3).

### Monkeys’ vigilance reactions towards the robot

Too few alarm calls have been emitted by the monkeys (Table 1) to perform statistical analyses but i) most of the callers were juveniles and ii) they produced snake and eagle alarm calls, instead of a ‘leopard alarm call’, which is usually used for a few species of carnivores^35^ as a predator deterrent^36^, suggesting that they did not consider the robot as a terrestrial predator. In fact, young vervet monkeys are more likely to produce alarm calls to a wider range of animals than adults who are more specific in producing alarm calls only to known predators^37^. Because the robot wore the same blue cap as the observers, provided food to the monkeys and that the observers walked close to the robot in the field, it is possible that the monkeys perceived the robot as a safety indicator, and even as a feeding opportunity. Moreover, this group of monkeys is very well habituated to humans with field experiments regularly conducted since 2011, which may have facilitated their curiosity towards novelty^38^. Although very few alarm calls were emitted, the monkeys displayed some vigilant behaviors such as ‘standing up bipedal’ and some self-centered behaviors such as ‘yawning’ and ‘self-scratching’. On the one hand, we found a significant effect of rank on the number of times individuals stood up bipedally (Zero-Inflated Poisson model: ZIP_1; Table SI_2) and on the number of self-centered behaviors (Zero-inflated negative binomial model: ZINB_1; Table SI_2). Higher-ranked individuals were 94 % more likely per unit of rank to stand-up bipedal than lower-ranked individuals (ZIP_1; p < 0.001; Table SI_2) and they were 95 % more likely per unit of rank to display self-centered behaviors than lower-ranked individuals (ZINB_1; p = 0.004; Table SI_2). Self-centered behaviors occur more frequently in stressful situations but the relation between rank and yawning and self-scratching appears unclear^39^. Here, it is possible that high rankers were torn between their motivation for food and their apprehension of the robot. Another potential explanation would be that observers missed these behaviors in low rankers because they were further away during the experiment, making these behaviors more difficult to detect in the bush. We also found a significant effect of sex on the number of times individuals stood up bipedally (ZIP_1; Table SI_2). Surprisingly, males were 52% less likely to stand up than females (p = 0.001; Table SI_2), while it has been reported that males are usually more vigilant than females who are more wary^40^.

On the other hand, we found no significant effect of age on the number of times the monkeys stood up bipedally (ZIP_1; p > 0.05; Table SI_2) and no significant effect of age and sex on the number of self-centered behaviors (ZINB_1; p > 0.05; Table SI_2).

### High rankers, females and juveniles monopolize the robot

We found a significant effect of rank on the number of scan points spent in close spatial proximity to the robot. Higher-ranked individuals were 88 % more likely per unit of rank to spend time close to the robot than lower-ranked individuals (ZINB_4; coefficient= -0.13; odds ratio = 0.88; p = 0.02; Figure 2). This effect can be linked to the significant effect of rank on the number of seconds spent eating food on the ground (ZINB_2; Table SI_2) and on the number of seconds spent eating food inside the box (ZINB_3; Table SI_2). Higher-ranked individuals were 92 % more likely per unit of rank to spend time eating food on the ground than lower-ranked individuals (p = 0.003; Table SI_2). Higher-ranked individuals were 84 % more likely per unit of rank to spend time eating food inside the box than lower-ranked individuals (p < 0.001; Table SI_2). These results are coherent with the fact that dominants commonly monopolize a food source, especially when it is a known and desirable food^41^. Individuals from the ‘G’ family were already among the first ones who foraged the most in a previous puzzle box experiment and a novel food test^42,43^, likely due to their high social status. It is also possible that high-ranking individuals recognized the box’s affordance as they experienced multiple box experiments in the past (reviewed in ^44^). A future study involving the ANYmal robot without a box is planned to investigate this effect.

**Figure 2.**
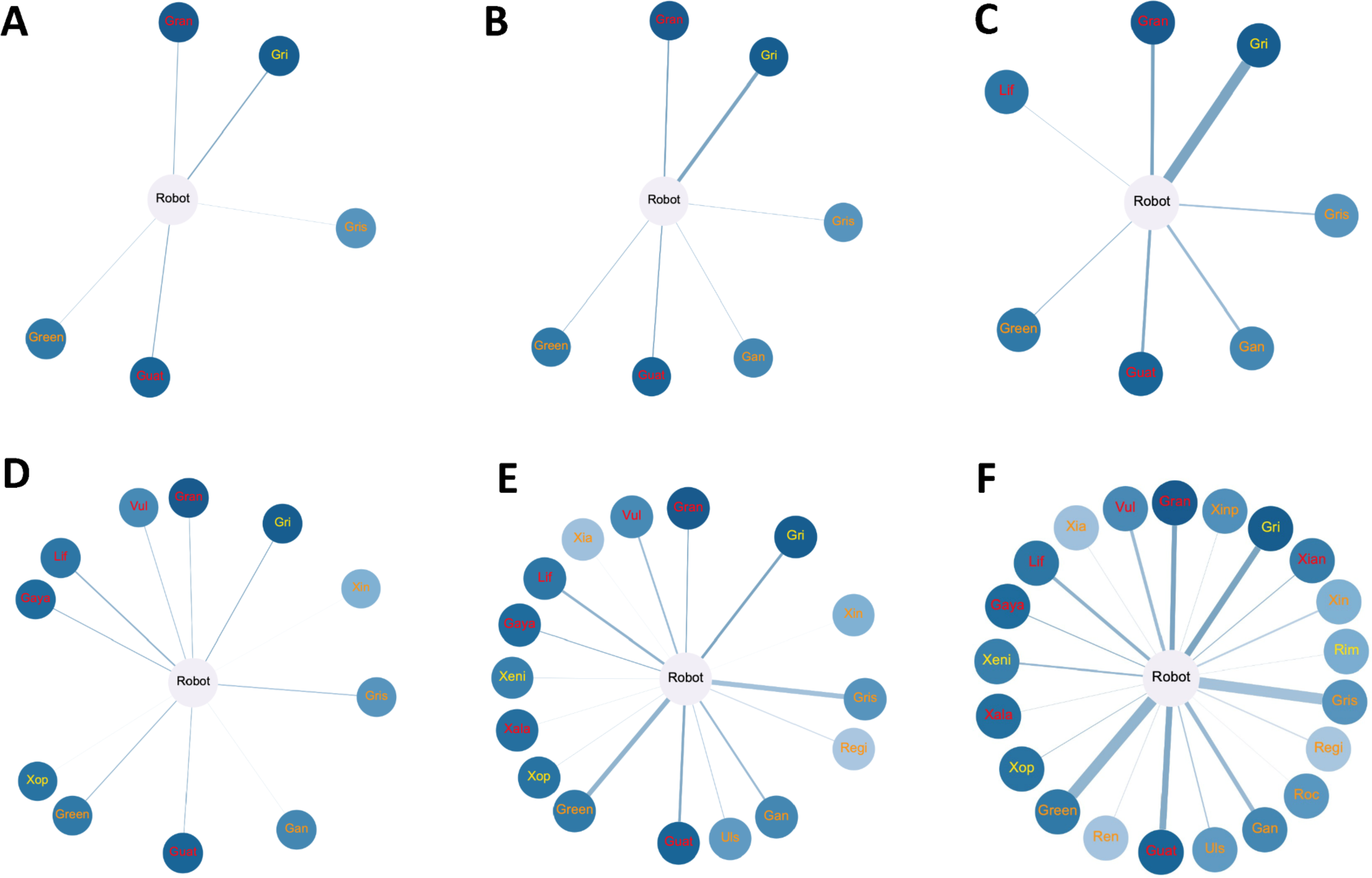
Spatial proximity networks around the robot across time. Cumulative close proximity networks (contact-arm length distances) A) on day 1; B) up to day 2 and C) up to day 6. Cumulative distant proximity networks (more than arm length distance and up to 10m) D) on day 1; E) up to day 2 and F) up to day 6. Each node represents an individual labelled by its name (three letters code for males, four letters code for females). Colours of the label name code for the age: adults are written in red; juveniles are written in orange and infants are written in yellow. The colour gradation of the nodes represents the hierarchical ranks: dark blue represents the higher-ranked individuals, while light blue represents lower-ranked individuals. Edges between the robot and the individuals represent the strength of association. The thicker the edge, the more the individual spent time in spatial proximity to the robot. Social networks were created with Gephi 0.10.1 software^45^.

We also found significant effects of sex and age on the number of seconds spent eating food on the ground (ZINB_2; Table SI_2) and on the number of seconds spent eating food inside the box (ZINB_3; Table SI_2). Females were 30 % more likely to eat food on the ground than males (ZINB_2; p = 0.01; Table SI_2). Females were 50% more likely to eat food inside the box than males (ZINB_3; p = 0.01; Table SI_2). This observed sex effect is driven by the fact that only one of the adult males and mostly females from the dominant ‘G’ matriline ate food around the robot and on the robot’s box.

Concerning age, Juveniles were 137,4 % more likely to eat food on the ground than adults (ZINB_2; p < 0.001; Table SI_2). Juveniles were 110,2 % more likely to eat food inside the box than adults (ZINB_3; p < 0.001; Table SI_2). These results are in accordance with previous studies reporting that juveniles were more likely to eat a novel food^46^, and they were more exploratory towards novel objects^47^, suggesting that they overcome neophobia faster and take more risks than adults^48^.

Finally, we found neither a significant effect of age and sex on the number of scans spent in close proximity to the robot (ZINB_4; p > 0.05; Figure 2) nor a significant effect of rank, age, and sex on the number of scans spent in more distant spatial proximity to the robot (ZINB_5; p > 0.05; Figure 2).

## Conclusion

Despite the technical challenges, we successfully introduced an ANYmal robot to a group of wild vervet monkeys. This study offers exciting opportunities for future animal-robot interaction studies in natural settings and for research questions such as: can monkeys learn from the robot and follow the robot’s food choice? Does a simple service make the robot accepted? Can knowledge from the robot be trusted? Can robots become group leaders? By implementing an artificial intelligence algorithm to visually identify individuals in the videos recorded by the robot through facial and body recognition, we could in future studies collect automatized social data to assess social networks.

## Methods

### Experimental model and subject details

One group of wild vervet monkeys (*Chlorocebus pygerythrus*), called ‘Noha’, took part in the study. NH was composed of 37 individuals (7 adult males; 10 adult females; 7 juveniles males; 7 juveniles females; 3 infants males; 3 infants females). Males were considered as adults once they dispersed, and females were considered as adults after they gave birth for the first time. Individuals that did not fulfil these criteria were considered as juveniles, with the exception of infants that were aged less than one year old. ‘Noha’ had been habituated to the presence of human observers since 2010. All individuals were identifiable thanks to portrait photographs and specific individual body and face features (scars, colours, shape etc.).

Ethics guidelines: Our study adhered to the “Guidelines for the use of animals in research” of Association for Study of Animal Behaviour and was approved by the relevant local authority, Ezemvelo KZN Wildlife, South Africa.

### Study site

The study was conducted within the INKAWU Vervet Project (IVP) in a 12000-hectares private game reserve: Mawana (28°00.327S, 031°12.348E) in the KwaZulu-Natal province, South Africa. The vegetation of the study site consisted in a savannah characterized by a mosaic of grasslands and clusters of trees of the typical savannah thornveld, bushveld and thicket patches. Mawana game reserve hosts various species of animals including elephants, giraffes, zebras, warthogs, and numerous species of antelopes. The common predators of vervet monkeys consist of hyenas, jackals, caracals, servals and several species of snakes and raptors.

### Hierarchy establishment

Agonistic interactions (e.g. stare, displacement, chase, hit, bite) were collected from January 2021 to October 2021 on all individuals of the group via *ad libitum* sampling method^49,50^ and food competition tests (i.e. corn provided to the whole groups from a plastic box). Data were collected by C.C and different observers from the IVP team. Before beginning data collection, observers had to pass an inter-observer reliability test with 80% of reliability for each data category between two observers. Data were collected on tablets (Vodacom Smart Tab 2) equipped with the Pendragon software version 8. Details about *ad libitum* data collection and hierarchy assessment have already been published in a previous paper^51^.

Individual hierarchical ranks were determined by the outcome of dyadic agonistic interactions recorded *ad libitum* and through food competition tests using the ‘EloRating’ package^52^ in R studio software version 2022.07.1^53^. Hierarchy of the group was significantly linear (h’ = 0.14; p = 0.013) and ranks were assessed by I&SI method^54^.

### Robot preparation

The ANYmal robot, a quadrupedal, sheep-sized robot originally developed by the Robotic Systems Lab at ETH Zürich, Switzerland, was adapted for this study in a game reserve. This adaptation involved adding a wide-angle camera module (Fig. 1A-ii), a long-range zoom camera (Fig. 1A-iii), and the food box (Fig. 1A-v). These modifications brought the robot’s weight to approximately 50 kg and allowed it to operate for about 50 minutes on a single battery charge.

To navigate the rough terrain of the game reserve, we employed a reinforcement learning (RL)-based locomotion controller^34,55^. The robot’s locomotion and balance were managed by a neural network controller, which processed proprioceptive measurements such as joint angles, velocity, and robot pose, alongside exteroceptive measurements from lidar sensors (Fig. 1A-iv). By integrating this multi-modal information, the neural network-based controller could generate real-time control commands for all 12 joints at a frequency of 50 Hz. We trained the controller with additional simulated disturbances for this experiment, enhancing the robot’s stability and safety even when monkeys jumped on or pushed it. The robot could traverse various rough terrains and cross a river within the reserve.

For continuous data collection and operation, we utilized the autonomy system developed by Team CERBERUS for the DARPA Subterranean Challenge^56^. This software was designed for the autonomous deployment of robots in cave-like environments, supporting both the remote control of the robot and its inspection head. Although we prepared a fully autonomous mode, it was not used during this study and we remote-controlled the robot around the monkeys for safety.

### Experiment

The field experiment consisted in introducing an ANYmal robot (Fig. 1) to the group of wild vervet monkeys. For this experiment, we added on the robot a blue cap (Fig. 1A-i) that is worn by researchers, and an open wooden box (filled with corn and apple slices) fixed on its back (Fig. 1). On the first day, the box was an opaque plastic box with an open lid, and it was fixed on the robot’s back. Due to unforeseen noise and vibration of the box when the robot was moving, on the second day, we changed the plastic box to a lighter wooden box of the same dimensions (20×20×15cm), without any lid, and kept it for the rest of the experiment. Although the robot can move autonomously, it was remotely controlled, at least 5m away, by J.L and S.Z for the purpose of the study. Note that we fixed the mobile head on the robot only for the last step of the experiment due to its fragility.

The experiment lasted one week from October 7^th^, 2021, to October 12^th^, 2021. Experiments took place once or several times a day, either at sunrise at the monkeys’ sleeping site or in the afternoon, depending on where the group was located and if it was easily accessible for the robot. C.C led the experiment with the help of M.A and one or two field assistants to directly identify the monkeys, along with J.L and S.Z who controlled the robot. Before the actual start of the experimental session, the robot approached the group of monkeys by walking in their direction and stopped a few meters away from them close to a potential refuge for the monkeys such as a tree. An experimental session started when the food was placed on the ground and/or inside the wooden box. From time to time during the trials, the researchers made some food calls, a lip-smacking call, to attract the monkeys. When the box was empty, E.v.d.W approached the box and baited it again. An experimental session ended when food was no longer available and/or when the group started moving away from the robot. During the experiment, all monkeys were free to approach the robot within the constraints of the social group dynamics. A total of 15 experimental sessions were run in ‘Noha’ for a total of 10h05min (Table 1; Table SI_1). The average duration of an experimental session was 40.33 minutes. Experiments were video recorded using a JVC camera (EverioR Quad Proof GZ-R430BE) to which C.C said aloud the identities of the individuals approaching the robot, eating the food on the ground and inside the box, being within 10m spatial proximity of the robot along with their behavioral reactions.

### Video analysis

All the video recordings were later analysed by C.C with VLC software version 3.0.16. Twenty percent of the video were also analysed by M.A and the inter-observer reliability was substantial (κ=0.70). During video analysis in slow motion or frame by frame, the following variables were encoded: the date, the exact time of each behavioral event and the identity of the actor and the recipient when relevant. Behavioral events were either considered as state events such as ‘eating food on the ground’ and ‘eating food inside the box’ for which we recorded the duration; or as point events to get the frequency of self-centered behaviors (yawn; self-scratch), agonistic behaviors (stare attack; head bob) and alarm calls towards the robot, vigilance behaviors (stand-up bipedal). Every minute, we used scan sampling^49^ to record the identity of individuals being in close spatial proximity to the robot (between in contact and arm-length distance) and in more distant spatial proximity to the robot (between more than arm length distance and 10m distance). We ended up with a total of 417 scans of spatial proximity over the whole experiment.

### Statistical analysis

Because our data sets comprised more than 50% of 0, we fitted Zero-Inflated regression models to our data (using the ‘glmmTMB’ package on R). We used the DHARMa package on R to assess all model diagnostics (dispersion test and zero inflation test on a GlmmTMB) and we visually checked the shape of Q-Q plots and residual deviation plots. Based on the models’ diagnostics and the comparison of the models AIC, we selected either Zero-Inflated Poisson regression models (ZIP) or Zero-Inflated Negative Binomial regression models (ZINB). For all the models, effect sizes are reported as odds ratio.

Models were run using the following R packages: ‘lme4’, ‘nlme’, MASS’, ‘pscl’ and ‘glmmTMB’ in R studio (version 2023.03.0+386).

#### Effects of sociodemographic factors on monkeys’ behaviors

We fitted a ZIP model (ZIP_1) to test for the effect of age, sex and rank on the number of vigilant behaviors (i.e. stand-up bipedal).

We fitted ZINB models to test for the effects of age, sex and rank on the number of self-centered behaviors displayed by monkeys (ZINB_1), and on the number of seconds they spent eating the food on the ground (ZINB_2) and inside the box (ZINB_3).

#### Effects of sociodemographic factors on spatial proximity to the robot

We fitted two ZINB models on observed spatial proximity data: ‘ZINB_4’ to test for the effect of sex, age and rank on the number of scan points spent in contact or within arm’s length distance to the robot and ‘ZINB_5’ to test for the effect of sex, age and rank on the number of scan points spent within more than arm length distance and up to 10m of distance to the robot. To deal with the non-independence of our spatial proximity data, we generated 2000 random data sets by permuting the sex, age and rank columns so values of number of scans spent in proximity to the robot are randomly allocated to individuals 2000 times. The ZINB models have been fitted for each of the 2000 permutations, generating a distribution of β estimates that we compared with the observed β. The null hypothesis was that the observed β coefficient is not different from the random set of β values. We rejected this hypothesis if the observed β value was lower/greater than 95% of the random values, meaning that the model estimate was significantly different from a random distribution.

## Acknowledgments

We thank the IVP onsite manager, Mike Henshall, and the whole IVP team for their help and support in the field, especially Aaron Mencia for his assistance in data collection. We are grateful to the van der Walt family for their permission to conduct the study on their land. We thank Frédéric Schütz and Loïc Brun for their advice and statistical help. This study has been funded by a collaborative grant from the Branco Weiss Fellowship—Society in Science granted to M.H and E.v.d.W. The INKAWU Vervet Project was funded during the time of this study by the Swiss National Science Foundation (PP00P3_198913) granted to E.v.d.W. At the time of writing, C.C was supported by the CNRS and by NeuroStra. This work of the Interdisciplinary Thematic Institute NeuroStra, as part of the ITI 2021-2028 program of the University of Strasbourg, CNRS, and Inserm, was supported by IdEx Unistra (ANR-10-IDEX-0002) under the framework of the French Program “Investments for the Future”. At the time of writing, E.v.d.W was supported by the European Research Council under the European Union’s Horizon 2020 research and innovation program for the ERC ‘KNOWLEDGE MOVES’ starting grant (grant agreement No. 949379).

## Authors contributions

The ANYmal robot has been prepared by M.H., J.L., and S.Z. and it has been controlled in the field by J.L, and S.Z. The initial conception of the project has been designed by M.H. and E.v.d.W. The experiment has been originally designed by C.C., J.L., S.Z., M.H, and E.v.d.W. M.M. made the food box and additional parts for robot. The experiment has been performed in the field by C.C., J.L., S.Z., M.A, and E.v.d.W. The video records of the experiment have been analyzed by C.C and M.A. The data have been analyzed by C.C. The figures have been prepared by C.C and J.L. The paper has been written by C.C and J.L. The funding has been acquired by M.H. and E.v.d.W. All authors reviewed and edited the draft.

## Research Ethics

The study was approved by the relevant local wildlife authority, Ezemvelo KZN Wildlife, South Africa (no reference number was provided). The study adhered to the “Guidelines for the use of animals in research” of the Association for the Study of Animal behaviour.

## Data availability

The datasets are made available at: https://doi.org/10.5281/zenodo.13309750

## Code availability

The R scripts for statistical analyses are available at: https://doi.org/10.5281/zenodo.13309750

