## Supplementary tables for "When monkeys meet an ANYmal robot in the wild"

| Step | Robot position | Food location | Date | Time | Session number | Experiment duration | Agonistic behaviours | Alarm calls | Eat food ground | Eat food box | First individual approaching the robot within 2m | First individual touching the robot | contact-arm length distance | > arm-length-10m distance |
|---|---|---|---|---|---|---|---|---|---|---|---|---|---|---|
| 1 | lay down | ground + box | 07/10/2021 | 11:40 | 1 | 24 | 0 | 0 | 0 | 0 | 0 | 0 | 0 | 0 |
| 1 | lay down | ground + box | 07/10/2021 | 16:31 | 2 | 19 | 0 | Gris, Prat, Upps | Gri; Gran; Gris; Green; Guat; Gan | 0 | Guat | 0 | Gri; Gran; Green | Guat, Gris, Lif; Gri; Gran; Green; Vul; Guat; Xin; Gaya |
| 1 | lay down | ground + box | 07/10/2021 | 17:08 | 3 | 12 | Gris | 0 | Guat; Gris; Gri | 0 | Guat | 0 | Guat; Gris; Gri | Lif; Gri; Gris; Xin; Vul |
| 1 | lay down | ground + box | 07/10/2021 | 17:27 | 4 | 37 | 0 | Guat | Gri; Gan; Green | 0 | Gri | 0 | 0 | Lif; Gri; Gaya; Gan |
| 1 | lay down | ground + box | 08/10/2021 | 05:32 | 5 | 53 | Gri | Xin; Uls | Guat; Gris; Gri; Green; Regi; Gran; Green; Vul; Lif; Uls; Xia | Gri; Gran; Green; Gris; Guat; Lif | Guat | 0 | Gris; Gri; Gran; Green; Guat; Lif | Guat; Gris; Gri; Green; Xeni; Xop; Vul; Regi; Gan; Uls; Xian; Xia |
| 1 | lay down | ground + box | 08/10/2021 | 06:30 | 6 | 26 | 0 | 0 | Gris; Gri; Gan; Green; Guat; Xia | 0 | Gan | 0 | 0 | Gris; Gan; Gri; Green; Guat; Xia; Xala |
| 2 | quadrupedal unstable body | ground + box | 08/10/2021 | 07:17 | 7 | 40 | Gri; Gris | 0 | Gris; Guat; Gri | 0 | Lif | 0 | 0 | Lif; Gris; Guat; Gri; Uls |
| 2 | quadrupedal unstable body | ground + box | 09/10/2021 | 05:39 | 8 | 57 | Gri | 0 | Green; Gris | Gan; Gris; Gri; Green; Guat; Gran | Gan | Gan | Gan; Gri; Gris; Guat; Green; Gran | Gan; Gris; Green; Gri; Lif; Guat; Gran; Xala; Vul; Xia; Regi |
| 3 | quadrupedal stable body | box | 09/10/2021 | 06:55 | 9 | 19 | 0 | 0 | 0 | 0 | Gri | Gri | 0 | 0 |
| 3 | quadrupedal stable body | box | 09/10/2021 | 07:45 | 10 | 38 | 0 | 0 | 0 | Gri; Green; Gran; Gris; Guat; Gan | Gan | Gri | Gri; Gran; Guat; Green; Gris; Gan | Gri; Green; Gris; Gan; Guat; Lif; Gran; Vul; |
| 4 | quadrupedal unstable body | box | 09/10/2021 | 16:51 | 11 | 26 | Gri | 0 | Guat | Gris; Guat; Gri; Gran; Gan | Gris | Gris | Guat; Gri; Gran; Gan | Gris; Green; Gran; Gri; Gan; Guat |
| 4 | quadrupedal unstable body | box | 10/10/2021 | 16:59 | 12 | 47 | Gri; Green | Guat | - | Gri; Gran; Gris; Gan; Guat | Gri | Gri | Gri; Gran; Gan; Guat | Gris; Gri; Guat; Green; Gran; Xin; Gan; Vul; Xinp |
| 5 | quadrupedal unstable body + turning head | box | 11/10/2021 | 08:41 | 13 | 66 | 0 | 0 | [Gris; Gan; Green] | Guat; Gri; Gran; Gris; Gan; Green | Guat | Guat | Guat; Gri; Gran; Gris; Gan | Guat; Lif; Gris; Green; Gran; Uls; Xia; Gan; Xeni; Rim; Xop; Xala; Xin; Xian |
| 5 | quadrupedal unstable body + turning head | box | 11/10/2021 | 16:25 | 14 | 60 | Gri | 0 | [Guat; Green; Xin; Gran] | Gan; Guat; Gri; Gris; Gran | Gan | Gan | Gan; Guat; Gri; Gris; Gran | Gan; Gran; Green; Gris; Guat; Gri; Xin; Regi |
| 5 | quadrupedal unstable body + turning head | box | 12/10/2021 | 05:40 | 15 | 81 | Gri; Green; Xin | Green; Xin | [Green; Gris; Lif] | Gan; Gri; Gran; Guat; Gris; Lif | Gan | Gan | Gan; Gri; Gran; Guat; Gris; Green; Lif | Gri; Xeni; Gris; Gran; Xin; Xian; Xop; Gan; Ren; Regi; Vul; Green |

Table SI_1. Summary table of the experiment by step; robot position; food position; date; time (start time of the experiment); session number; experiment duration (in minutes: seconds); list of individuals who emitted agonistic behaviours towards the robot; ist of individuals who emitted alarm calls in response to the robot; list of individuals eating food on the ground; [Names into brackets represent the individuals that ate food leftovers on the ground]; list of individuals eating food inside the box on the robot’s back; list of individuals present in contact and within arm length distance to the robot during scans; list of individuals present within more than arm length distance to 10m to the robot during scans.

| Model label | Model | Parameter | Coefficient | Odds ratio | Standard error | P value |
|---|---|---|---|---|---|---|
| ZIP_1 | log | Intercept | 2.94 | 18.92 | 0.18 | < 0.0001 |
| | | sex_Male | -0.73 | 0.48 | 0.23 | 0.001 |
| | | Age_Juvenile | 0.29 | 1.34 | 0.26 | 0.26 |
| | | Rank | -0.06 | 0.94 | 0.01 | < 0.001 |
| | logit | Intercept | 0.44 | 1.55 | 0.83 | 0.60 |
| | | sex_Male | -1.19 | 0.30 | 0.84 | 0.16 |
| | | Age_Juvenile | -1.42 | 0.24 | 0.90 | 0.12 |
| | | Rank | 0.09 | 1.09 | 0.05 | 0.05 |
| ZINB_1 | log | Intercept | 2.92 | 18.54 | 0.33 | < 0.001 |
| | | sex_Male | 0.04 | 1.04 | 0.32 | 0.89 |
| | | Age_Juvenile | 0.70 | 2.01 | 0.39 | 0.08 |
| | | Rank | -0.05 | 0.95 | 0.02 | 0.02 |
| | logit | Intercept | 0.37 | 1.45 | 0.77 | 0.63 |
| | | sex_Male | -0.64 | 0.53 | 0.73 | 0.39 |
| | | Age_Juvenile | -1.22 | 0.30 | 0.79 | 0.12 |
| | | Rank | 0.05 | 1.05 | 0.04 | 0.17 |
| ZINB_2 | log | Intercept | 5.41 | 223.63 | 0.44 | < 0.001 |
| | | sex_Male | -1.20 | 0.30 | 0.47 | 0.01 |
| | | Age_Juvenile | 2.62 | 13.74 | 0.61 | < 0.001 |
| | | Rank | -0.08 | 0.92 | 0.03 | 0.003 |
| | logit | Intercept | 0.48 | 1.62 | 0.79 | 0.55 |
| | | sex_Male | -0.38 | 0.68 | 0.76 | 0.62 |
| | | Age_Juvenile | -1.18 | 0.31 | 0.83 | 0.16 |
| | | Rank | 0.06 | 1.06 | 0.04 | 0.13 |
| ZINB_3 | log | Intercept | 7.64 | 2079.7 | 0.23 | < 0.001 |
| | | sex_Male | -0.70 | 0.50 | 0.29 | 0.01 |
| | | Age_Juvenile | 2.40 | 11.02 | 0.33 | < 0.001 |
| | | Rank | -0.17 | 0.84 | 0.02 | < 0.001 |
| | logit | Intercept | -0.07 | 0.93 | -0.08 | 0.94 |
| | | sex_Male | -0.87 | 0.42 | -0.79 | 0.43 |
| | | Age_Juvenile | -0.60 | 0.55 | -0.55 | 0.58 |
| | | Rank | 0.17 | 1.19 | 2.27 | 0.02 |

Table SI_2. Model outputs for Zero-Inflated Poisson regression model (ZIP_1) and Zero-Inflated Negative Binomial regression models (ZINB_2, ZINB_2 and ZINB_3).
